# Apparent oxygen half saturation constant for nitrifiers: genus specific, inherent physiological property, or artefact of colony morphology?

**DOI:** 10.1101/289645

**Authors:** Yingyu Law, Artur Matysik, Xueming Chen, Sara Swa Thi, Thi Quynh Ngoc Nguyen, Guang Lei Qiu, Gayathri Natarajan, Rohan B.H. Williams, Bing-Jie Ni, Thomas William Seviour, Stefan Wuertz

## Abstract

We report that a single *Nitrospira* sublineage I OTU performs nitrite oxidation in several full-scale domestic wastewater treatment plants (WWTPs) in the tropics (29-31 °C). Contrary to the prevailing theory for the relationship between nitrite oxidizing bacteria (NOB) and ammonia oxidizing bacteria (AOB), members of the *Nitrospira* sublineage I OTU had an apparent half saturation coefficient, *K*s_(app)_ lower than that of the full-scale domestic activated sludge cohabitant AOB (0.09 ± 0.02 g O2 m^−3^ versus 0.3 ± 0.03 g O2 m^−3^). Paradoxically, NOB may thus thrive under conditions of low oxygen supply. Low dissolved oxygen (DO) conditions could enrich for and high aeration inhibit the NOB in a long-term lab-scale reactor. The relative abundance of *Nitrospira* gradually decreased with increasing DO until it was washed out. Nitritation was sustained even after the DO was lowered subsequently. Based on 3D-fluorescence *in situ* hybridization (FISH) image analysis, the morphologies of AOB and NOB microcolonies responded to DO levels in accordance with their apparent oxygen half saturation constant *K*s_(app)_. When exposed to the same oxygenation level, NOB formed densely packed spherical clusters with a low surface area-to-volume ratio compared to the *Nitrosomonas*-like AOB clusters, which maintained a porous and non-spherical morphology. Microcolony morphology is thus a way for AOB and NOB to regulate oxygen exposure and sustain the mutualistic interaction. However, short-term high DO exposure can select for AOB and against NOB in full-scale domestic WWTPs and such population dynamics depend on which specific AOB and NOB species predominate under given environmental conditions.

## Introduction

Nitrification activates inert reduced inorganic nitrogen (i.e. ammonium) in the presence of oxygen to its oxidized form nitrate via nitrite. It is a crucial step in global biogeochemical nitrogen cycling as well as in biological wastewater treatment (Ward 2011). Ammonia and nitrite oxidation is catalyzed either in a two-step process by phylogenetically distinct ammonia-oxidizing bacteria (AOB) or archaea (AOA), and nitrite-oxidizing bacteria (NOB), respectively (Bock and Wagner 2001, Könneke et al 2005) or directly by complete ammonia oxidizers (comammox) (Daims et al 2015, Van Kessel et al 2015). In the two-step process, AOB and NOB are often in close proximity to one and another in order to finely balance production and consumption of the potentially toxic nitrite (Gieseke et al 2003, Matsumoto et al 2010, Stein and Arp 1998, Teske et al 1994). NOB are dependent on AOB for their electron donor, yet they also compete with AOB for oxygen as a terminal electron acceptor, particularly under oxygen limiting conditions (Juretschko et al 1998b, Sliekers et al 2005). As a consequence of this tight-knit interaction between AOB and NOB, changes in the activity and relative abundance of AOB in response to environmental perturbations could significantly impact the stability of NOB activity (Knapp and Graham 2007). For example, in wastewater treatment systems, the oxygen competition dynamic between AOB and NOB is utilized to repress the growth of NOB, enabling stable interaction between AOB and anammox (anaerobic ammonium oxidizing) bacteria (Lackner et al 2014). Anammox bacteria can convert ammonium directly to dinitrogen gas using nitrite as an oxidant (Strous et al 1998) and when coupled with partial ammonium conversion exclusively to nitrite (i.e. partial nitritation) by AOB, complete nitrogen removal can be achieved with far less energy expenditure (Siegrist et al 2008).

Relative half saturation constant (*K*s) and affinities for oxygen (maximum specific growth rate per half-saturation constant, *µ*_max_/*K*s) can be used to inform and control the competition dynamics between AOB and NOB (Wett et al 2013). While there have been numerous studies that have investigated the interactions between AOB and NOB, mainly in aquatic systems or biofilm- and activated sludge-based wastewater treatment systems (Gieseke et al 2003, Jubany et al 2008, Ke et al 2013, Wiesmann 1994), there is still uncertainty about their relative *K*s for oxygen. It is generally accepted that AOB have a higher affinity for oxygen (i.e. a lower *K*s for oxygen) than NOB. Consequently most full-scale partial-nitritation installations treating high-strength wastewater (total nitrogen > 100 mg N/L), for example, adopt low oxygen set points of less than 0.35 mg O_2_/L to curb NOB activity (Lackner et al 2014). Such observations are based predominately on studies in high-strength wastewater treatment processes with high ammonia concentrations (Jubany et al 2008, Wiesmann 1994). However, there are other studies which suggest that NOB have lower *K*s than AOB based mainly on results from activity tests under domestic wastewater conditions (total nitrogen between 20 and 70 mg N/L) (Malovanyy et al 2015, Regmi et al 2014, Sliekers et al 2005).

The discrepancy observed is likely influenced by phylogenetic variability of AOB and NOB in different systems. Members of the genus *Nitrosomonas* are among the most frequently detected AOB in wastewater treatment plants (WWTPs) (Purkhold et al 2000), but different treatment plants can be dominated by either single or multiple *Nitrosomonas*-like AOB populations (Daims et al 2001b, Juretschko et al 1998a). In the case of NOB, *Nitrospira* and *Nitrobacter* have also been reported to co-exist in laboratory-scale reactors and full-scale treatment plants where nitrite concentrations fluctuate within operational cycles, which allow for niche differentiation (Coskuner and Curtis 2002, Daims et al 2001b, Kim and Kim 2006). The genus *Nitrospira* that predominates under domestic wastewater conditions generally has a lower *K*s for oxygen and nitrite than *Nitrobacter*, which occurs in nitrogen rich systems (Daims et al 2016, Regmi et al 2014). In fact, full genome analysis of *Candidatus* Nitrospira defluvii enriched from a sewage treatment plant revealed the lack of basic defence mechanisms against oxidative stress suggesting that their nitrite oxidising activity could potentially be inhibited at high DO concentration (Lücker et al 2010). In addition to nitrifying community diversity, differences in the AOB to NOB ratio can also significantly impact their interaction(Yao and Peng 2017). Such variations in relative abundance and physiological properties suggest that the relative *K*s for oxygen will depend on the AOB and NOB populations present, and may in fact be system specific. Therefore, the same inhibitory conditions for NOB in high wastewater strength processes may not work under domestic wastewater conditions.

The measurement of *K*s and *µ*_max_ for oxygen is also impacted by the physical properties of the microbial aggregates (Martins et al 2004, Picioreanu et al 2016). In activated sludge wastewater treatment systems, nitrifiers grow in microcolonies within flocs (Daims et al 2001b, Mobarry et al 1996, Wagner et al 1995) which affects the mass transfer rate of substrate to the cells. Picioreanu *et al*. (2016) showed by 3D modelling that the reversal of *K*s (i.e. NOB lower than AOB) is an artifact of the generally larger average microcolony sizes of AOB compared to NOB coupled with the higher oxygen yield of AOB over NOB, with the former having greater influence on the *K*s for oxygen. DO is consumed more quickly in AOB clusters, resulting in DO limitation and lower measured activity. The authors concluded that incorporating a description of the physical structure of the AOB and NOB clusters is crucial to understanding oxygen competition between AOB and NOB.

The aim of this study was, therefore, to resolve the contributions of different genera and microcolony morphologies to the oxygen half saturation constant (*K*s) of NOB and AOB. Deconvoluting the contributions of genera and microcolony morphologies to *K*s was achieved by combining AOB and NOB microbial community characterisation in activated sludge with the determination of their respective apparent oxygen half saturation constant, *K*s_(app)_ and physical properties of the AOB and NOB clusters using 3D-fluorescence *in situ* hybridization (FISH) and image analysis. Given that substrate diffusion into the microcolonies may be rate limiting, we consider the measured *K*s_(app)_ values as an upper estimate for the half-saturation constant. In addition, the effect of varying oxygenation levels on AOB and NOB activity was also investigated in a long term lab-scale reactor. The findings suggest that cluster shape and size are a consequence of rather than the cause of different *K*s_(app)_ which may serve as a mechanism that enables AOB and NOB with distinct preference for oxygen to coexist within the same ecological niche at a certain range of DO concentration. This also informs competition dynamics between two key players in nitrification and can lead to improved DO control strategies for achieving nitritation for more cost effective wastewater treatment.

## Materials and Methods

### Sampling of activated sludge from major WWTPs treating domestic wastewater

Activated sludge was sampled from three main WWTPs (Plants 1, 2 and 3) collectively treating 90% of the total domestic wastewater loading in Singapore. Three sampling events were carried out for each plant between December 2016 and February 2017. On each sampling event, liquid samples were collected from the influent and effluent of the activated sludge treatment unit. The collected liquid samples were filtered with 0.22-µm disposable sterile filters and analyzed for ammonium, nitrate, nitrite and orthophosphate. Non-filtered liquid samples collected from the influent were immediately acidified with sulfuric acid and analyzed for total chemical oxygen demand (TCOD). Samples for DNA extraction and fluorescence *in situ* hybridization (FISH) were collected from the mid aerobic zone. Samples for DNA extraction were snap-frozen in liquid nitrogen and stored at −80 °C until extraction. Samples for FISH were immediately fixed in 4% paraformaldehyde and were washed twice with 1% phosphate-buffered saline solution after 2 h and stored at −20 °C in a 50:50 mixture of 1% phosphate-buffered saline solution and 100% ethanol.

### Long-term lab-scale reactor start-up and operation

A sequencing batch reactor (SBR) with a working volume of 4 L was seeded with returned activated sludge (RAS) from Plant 3 that receives domestic wastewater at a rate of 800,000 m^3^/day. Wastewater was collected from the plant once a week and stored in a chiller at 4 °C as feed for the SBR. The nutrient content of the wastewater was analysed for every batch of wastewater and is summarised in Table S1 (Supporting Information). The SBR was operated to achieve partial nitritation mode (50% conversion of ammonium to nitrite, with the remaining ammonium unconverted) for the first 150 days of operation and switched to nitritation (complete conversion of ammonium to nitrite) mode thereafter. Anoxic phases for denitrification were also included in the SBR operation to exhaust the organics in the wastewater as well as to restore the alkalinity in the reactor. Therefore, pH was not specifically controlled in the reactor but stabilised between 6.3 and 7.0. The effect of DO concentration on nitrite accumulation was investigated by varying the DO setpoint between 0.5 and 5.5 mg O_2_/L. In addition, the effect on the length of aeration period on the AOB and NOB activities were also investigated by switching the SBR operation from two to three feeding phases while maintaining the overall wastewater loading rate with a total SBR cycle time of eight hours. When the SBR was operated in the cycle with two feeding phases, 1.5 L of wastewater was fed into the reactor over 75 min of slow feeding and each feeding phase was followed by 60 min of anoxic phase and 60 min of aerobic phase. The whole cycle ended with 90 min of settling and decanting. When switched to the cycle with three feeding phases, the SBR was fed with 1.0 L of wastewater over each 50 min feeding. The following anoxic phases were shortened to 30 min and aerobic phases were shortened to 18-40 min depending on the DO set point. The length of the settling and decant phase was adjusted accordingly to maintain the overall cycle at eight hours, thus resulting in a hydraulic retention time (HRT) of 10.7 h. A heating jacket was connected to maintain the SBR temperature at 31.0 ± 0.5 °C, typical temperature of domestic wastewater in Singapore. DO and pH were continuously monitored using Mettler Toledo-InPro 3250i pH sensor and Mettler Toledo InPro6050 DO sensor, respectively. The solid retention time (SRT) was not controlled for the first 160 days of reactor operation and was estimated to be 14.4 ± 2.0 days due to the inevitable sludge loss with the effluent. When stable partial nitritation was achieved, the SBR was switched to nitritation mode from day 151 onwards by increasing the length of the aerobic phase to investigate whether NOB will recur with extended aeration. In addition, the SRT was controlled at approximately 8 days from day 161 onwards, based on measured total suspended solids (TSS) of the mixed liquor in the SBR and effluent TSS. Each operational change in DO set point, aeration cycle length (either during the switch from two to three feeding phases or from partial nitritation to nitritation mode) and solid retention time was implemented individually and separated as different experimental phases (I to X) as summarised in Table 2. Samples were collected periodically at the end of the cycle and filtered immediately with 0.2 µm filters.

**Table 1.** The key operational condition, effluent quality and corresponding relative abundance of AOB and NOB in three major wastewater treatment plants treating domestic waste water (values are averages and standard error of the mean from three sampling events).

|  | Plant 1 | Plant 2 | Plant 3 |
| --- | --- | --- | --- |
| Process configuration | Pre-denitrification/Anaerobic / Aerobic/ MBR | Modified Ludzack Ettinger | Anoxic/Oxic Step Feeding |
| Dissolved Oxygen (mg O<sub>2</sub>/L) | 1.6±0.6 | 0.8±0.2 | 1.2±0.2 |
| pH | 7.7-8.3 | 6.9-7.7 | 6.4-7.0 |
| Solid retention time (day) | 15-18 | 4-6 | 4-6 |
| Effluent ammonium (mg N/L) | 0.2±0.2 | 3.1±1.5 | 2.4±1.0 |
| Effluent nitrate (mg N/L) | 8.1±0.8 | 7.5±0.9 | 10.6±8.0 |
| Effluent nitrite (mg N/L) | 0 | 0.7±0.2 | 0.3 |
| Effluent phosphate (mg P/L) | 0.8±0.8 | 0 | 1.6±0.7 |
| Effluent TCOD (mg/L) | 17±2 | 17.9±1.5 | 11.3±0.9 |
| Mixed liquor suspended solids (g/L) | 3.16±0.07 | 2.06±0.14 | 2.44±0.01 |
| Relative abundance of *g_Nitrospira* (%) | 0.83±0.05 | 1.67±0.38 | 1.81±0.30 |
| Relative abundance of *f_Nitrosomonadaceae* (%) | 1.79±0.12 | 0.64±0.05 | 0.33±0.09 |

**Table 2.**
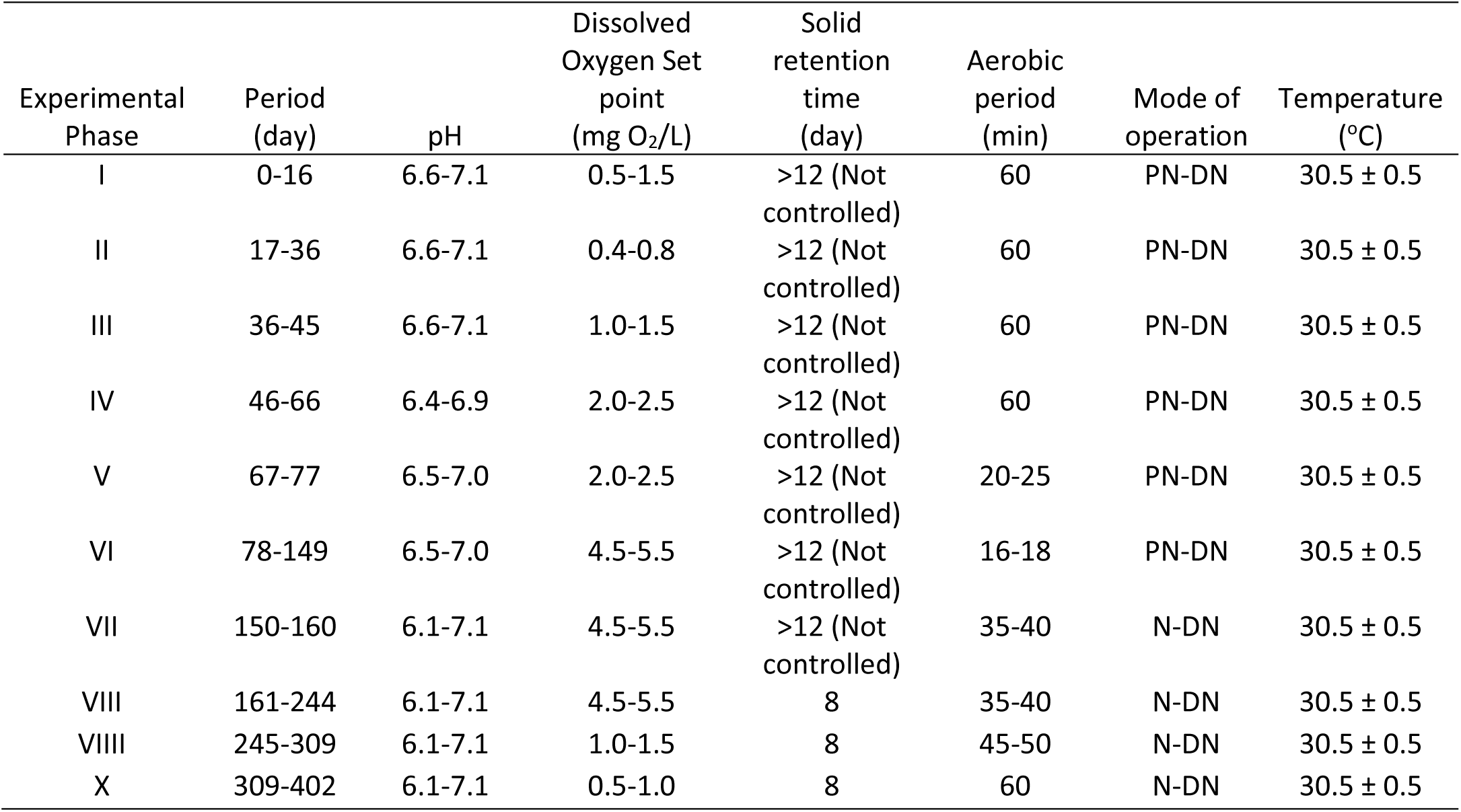
Operational parameters at different experimental stages of the lab-scale partial nitritation (PN)/ nitritation (N)-denitritation (DN) sequencing batch reactor.

### Batch activity experiments & analytical procedures

Three types of activity batch tests were conducted using Plant 3 returned activated sludge (RAS): (1) AOB activity test, (2) NOB activity test and (3) simultaneous AOB and NOB activity tests. Given that AOB and NOB microcolonies are found to cluster in close proximity to one and another the simultaneous AOB and NOB activity tests are important to elucidate how oxygen consumption would impact oxygen availability and hence each other’s activity (Picioreanu et al 2016). All three activity tests were conducted at DO concentrations of 0.5 ± 0.1, 1.5 ± 0.1, 2.5 ± 0.1, 5.5 ± 0.1 and 6.5 ± 0.1 mg O_2_/L. Additional DO concentrations of 0.2 ± 0.1 and 0.3 ± 0.1 mg O_2_/L were carried out for the NOB activity test and the simultaneous AOB and NOB activity test, respectively. DO concentration was manually controlled in all batch tests using a gas mixture of N2 and air independently connected to two rotameters. Fresh sludge was collected every 48 h and diluted with effluent from the lab-scale SBR at a 1:1 ratio before each batch experiment. When steady nitritation activity was achieved in the lab-scale SBR, sludge was also collected to perform AOB activity batch test at the aforementioned DO concentrations. Ammonium or nitrite was added at the start of the AOB or nitrite oxidising batch activity tests to an initial concentration of 25 mg N/L and 15 mg N/L, respectively. Both ammonium and nitrite were provided to initiate the simultaneous AOB and NOB activity tests. All activity tests were conducted in triplicate with the pH controlled at 7.0 ± 0.1 with addition of 0.5 M sodium bicarbonate solution. Nutrient samples were collected every 10-20 min and filtered immediately with 0.2 µm filters. Mixed liquor samples were collected for suspended solids (MLSS) and volatile suspended solids (MLVSS) analyses at the start of every batch experiment.

Filtered samples were analysed for ammonium, nitrite, nitrate, orthophosphate and soluble COD. Ammonium and COD were measured using Hach® kits whereas nitrate, nitrite and orthophosphate were analysed using ion chromatography (Prominence, Shimadzu). Total Kjeldahl Nitrogen (TKN) was measured using a total nitrogen measuring unit (Shimadzu). MLSS and MLVSS were analyzed according to the standard methods (APHA 2005).

### Estimation of AOB and NOB associated kinetics

A Monod-based model with well-established biokinetics of AOB and NOB was applied (Wiesmann 1994) to gain further insight into the microbial interactions between AOB and NOB involved in this work. Table S2 in the Supporting Information summarizes the stoichiometrics and kinetics of the model, while Table S3 lists the definitions, values, units and sources of all parameters used in the model. In this work, we particularly evaluated the maximum growth rate of AOB (μ_AOB_) and NOB (μ_NOB_) and the apparent oxygen half saturation coefficient for AOB (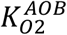) and NOB (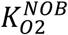) considering their key roles in regulating the microbial competition between AOB and NOB, while the literature values for affinity constants of ammonium and nitrite were used due to the excessive supply of ammonium (20-25 mg N/L) or nitrite (15 mg N/L) applied in the activity batch tests in comparison to the reported relatively low affinity constants of ammonium and nitrite as shown in Table S2.

Specifically, the NOB biokinetics (i.e., μ_NOB_ and 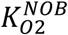) were first calibrated and validated using the NOB activity batch tests conducted on the Plant 3 sludge fed with nitrite at various controlled DO levels (0.2, 0.5, 1.5, 2.5, 5.5 and 6.5 mg/L). On top of the NOB biokinetics obtained, the calibration and validation of the AOB biokinetics (i.e., μ_AOB_ and 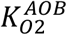) was then carried out using AOB activity batch tests conducted on Plant 3 sludge fed with ammonium at different DO levels of 0.5, 1.5, 2.5, 5.5 and 6.5 mg/L. The NOB and AOB biokinetics was also validated with simultaneous AOB and NOB activity batch tests data. The obtained biokinetics of AOB and NOB was further applied to verify the dominance of AOB over NOB after enrichment using batch tests conducted on the lab enriched sludge fed with ammonium at the aforementioned DO levels. Biomass concentrations used in the model evaluation process were set based on the MLVSS measurements in combination with the microbial abundance analysis detailed in the following section. The model was implemented on the software AQUASIM 2.1d (Reichert 1998) and configured according to the specific conditions of the batch tests as described in the earlier section.

#### Microbial Community Characterisation

Biomass samples collected periodically from the lab-scale reactor and from the wastewater treatment plants for DNA extraction were stored at −80°C. The DNA was extracted based on the FastDNATM 2 mL SPIN Kit for Soil (MP Biomedi-cals, USA) optimised for DNA extraction from activated sludge (Albertsen et al 2015). 16S rRNA gene amplicon sequencing was conducted by DNASense (http://dnasense.com/) to determine the microbial community structure of the activated sludge. Bacterial primers 27F (AGAGTTTGATCCTGGCTCAG, Lane 1991) and 534R (ATTACCGCGGCTGCTGG, (Muyzer et al. 1993)) were used to amplify an approximately 500 bp DNA fragment of the 16S rRNA gene (variable V1 to V3 regions). Amplification of PCR was done using the following conditions: 2 min of 95°C, 20 sec for 30 cycles of 95°C, 30 sec of 56°C, 60 sec of 72°C, and a 5 min elongation step at 72°C. This amplification utilized 1X Platinum® High Fidelity buffer, 2 mU Platinum® Taq DNA Polymerase High Fidelity, 400 pM dNTP, 1.5 mM MgSO4, 5 uM V1-V3 adaptor mix (barcoded), and 10 ng of template DNA. Purification of PCR products was done using the Agencourt AmpureXP (Beckman Coultier Inc., U.S.A.) with 1.8 bead solution/PCR solution ratio. The QuantIT HS kit (Life Technologies, USA) was used to quantify the DNA concentration. Using Illumina MiSeq (Illumina Inc., USA), barcoded amplicons, which were pooled in equimolar amounts, were paired-end sequenced (2×250 bp).

The output from the MiSeq (Illumina Inc., San Diego, California, USA) was de-multiplexed from the amplicon libraries in FASTQ-format for each sample in the composite library. Pre-processing of all amplicon libraries was performed according to Albertsen *et al*. (Albertsen et al 2015), and all sequenced sample libraries were subsampled to 10,000 raw reads. Taxonomy was assigned using MiDAS v.1.20 (McIlroy et al 2015) with 97% clustering identity.

### Fluorescence in situ Hybridisation (FISH) and image processing

The method described by Daims et al. (Daims et al 2001a) was used to prepare biomass samples for FISH analysis. The following probes were used: NSO1225 and NSO190, specific for ammonia oxidizing betaproteobacteria; Ntspa662 targeting all *Nitrospira*; and EUB-mix (EUB338, EUB338-II, and EUB338-III), covering most bacteria. All probes were either labeled with indocarboncyanine (Cy3 or Cy5) or 6-carboxyflorescein (6-FAM). Z-stacks of FISH-probed samples were acquired using a Zeiss LSM 780 inverted confocal microscope equipped with 100x/NA1.4 oil immersion objective (Carl Zeiss, Jena, Germany) and pre-processed using FIJI (background subtraction, 3D median) (Schindelin et al 2012). DAIME software was used for 3D visualization, segmentation and quantification of bacterial clusters (Daims et al 2006). Surface, volume, surface-to-volume ratio and shape were measured for every recognized cluster in respective AOB and NOB channels for 20 distinct regions-of-interest (ROIs). Unpaired nonparametric Mann-Whitney U test was conducted to assess differences between AOB and NOB clusters.

## Results

### Diversity of nitrifying communities in major WWTPs

A single *Nitrospira* sublineage I OTU was the predominant NOB member in all three treatment plants sampled (Figure 1), despite differences in process design and operational conditions. In contrast, up to 18 OTUs in the family Nitrosomonadaceae were detected by 16S rRNA gene amplicon sequencing analysis, of which only eight were assigned to the genus *Nitrosomonas* (Figure 1). Other known ammonia oxidizing and nitrite oxidizing taxa were not detected. The collective relative abundance of OTUs of AOB was lower than that of NOB in Plants 2 and 3, with ratios of NOB to AOB of 2.6 and 5.5, respectively (Figure 1 and Table 1). The opposite was observed for Plant 1 where AOB were present at a higher abundance relative to NOB with a ratio of NOB to AOB of 0.5. Apart from the difference in plant design, Plant 1 had the highest DO concentration in the aerobic zone and the longest SRT compared to Plants 2 and 3 (Table 1). The lower relative abundance of AOB in these plants is also reflected in the ammonium removal efficiency with a higher residual ammonium concentration compared to that of Plant 1 (Table 1). This increase in AOB:NOB ratio under higher DO condition in a full-scale WWTP is consistent with the observation that NOB have higher relative oxygen affinities. High DO could also be potentially toxic to *Nitrospira*. Thus, to confirm that DO can be used to select against NOB, sludge from Plant 3 (with a high NOB:AOB ratio) was transferred to a lab-scale reactor, exposed to a range of DO conditions, and the AOB to NOB competition dynamics were observed.

**Figure 1.**
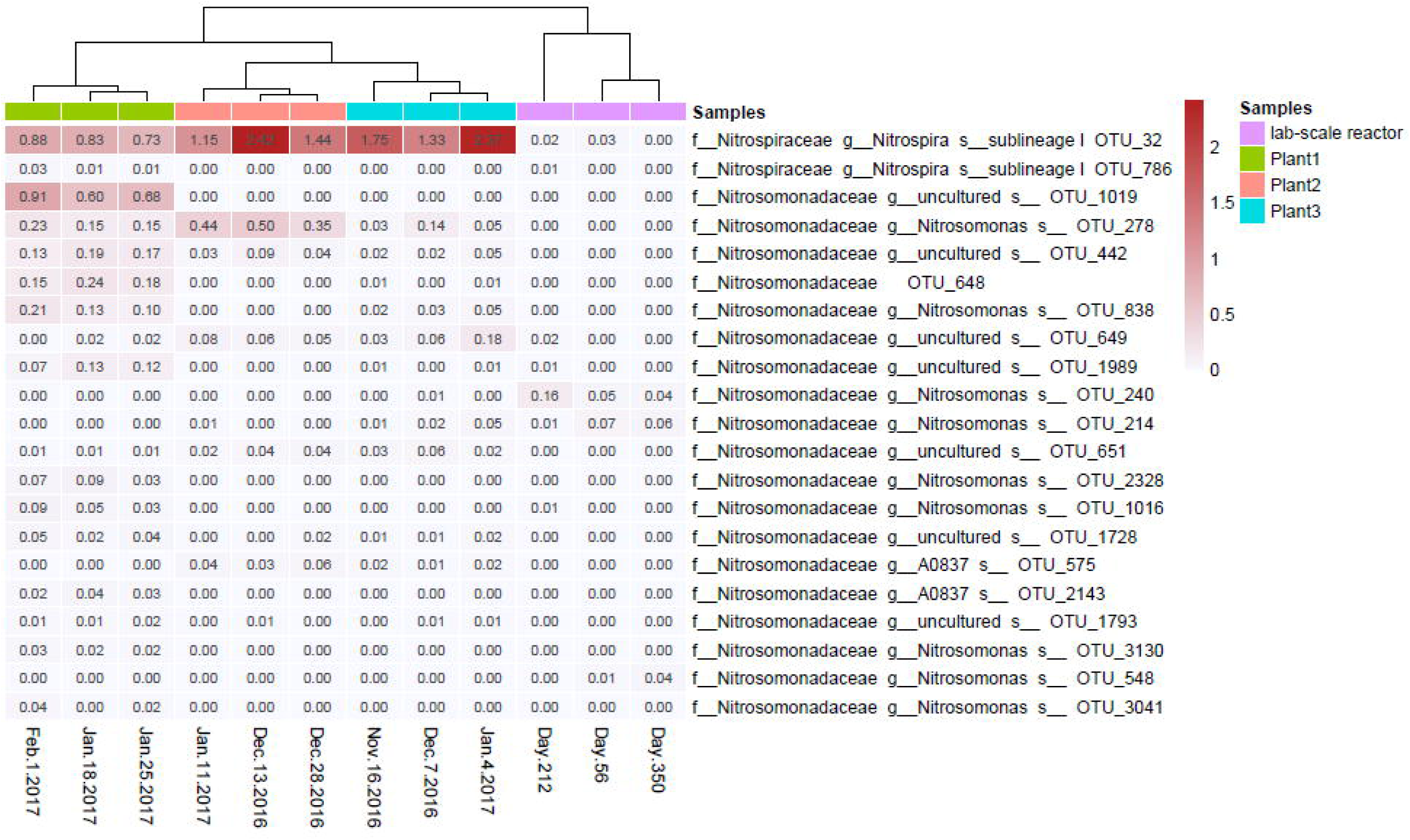
Ammonia oxidizing and nitrite oxidizing bacteria detectable by16S rRNA gene amplicon sequencing in three major wastewater treatment plants in Singapore and the lab-scale partial nitritation/ nitritation-denitritation reactor during different experimental phases (see Table 2 and Figure 2). The scale on the heat map denotes relative abundance of each taxon in %.

### Dynamics of microbial community structure in response to varying DO set points

The DO concentration and length of aeration phase were systematically varied in each experimental phase to investigate DO selection against NOB in a lab-scale reactor (Table 2). Consistent with what was observed in the full-scale plant, only a single *Nitrospira* sublineage I OTU was detected whereas multiple OTUs were annotated to Nitrosomonadaceae throughout the experiment. In experimental phases I, II and III, all of the ammonium oxidised by AOB was converted to nitrate by NOB when DO set points of < 1.5 mg O_2_/L were applied (Table 2 and Figure 2A). *Nitrospira* sublineage I had a much higher relative abundance than AOB during these initial experimental phases (Figure 2B). The increase in DO set point to 2.0 – 2.5 mg O_2_/L resulted in a slight increase in the nitrite:NOx (sum of nitrite and nitrate in the effluent) ratio in experimental phase IV and a corresponding reduction in the relative abundance of *Nitrospira* from 3.7% to 2.1%, albeit it was still more abundant than AOB (Figure 2B). When the aeration phase was shortened from 60 min to 20 – 35 min while maintaining the DO set point at 2.0 – 2.5 mg O_2_/L (Table 2), no apparent change was observed in both the relative abundance of *Nitrospira* and nitrite production. *Nitrospira* gradually decreased to below the detection limit in experimental phase VI, when the DO set point was further increased to 4.5 – 5.5 mg O_2_/L, and partial nitritation was achieved with a concomitant increase in final effluent nitrite concentration and the overall nitrite:NOx ratio (Figure 2A and B). Nitrite accumulation was sustained with subsequent changes in operational conditions, including 1) extension of the aerobic period to achieve complete conversion of ammonium to nitrite (nitritation) in experimental phase VII; 2) decrease in SRT in experimental phase VIII; and 3) return of the DO concentration back to 1.0 – 1.5 mg O_2_/L in experimental phase VIIII and 0.5 – 1.0 mg O_2_/L in experimental phase X (Figure 2A and B, Table 2).

**Figure 2.**
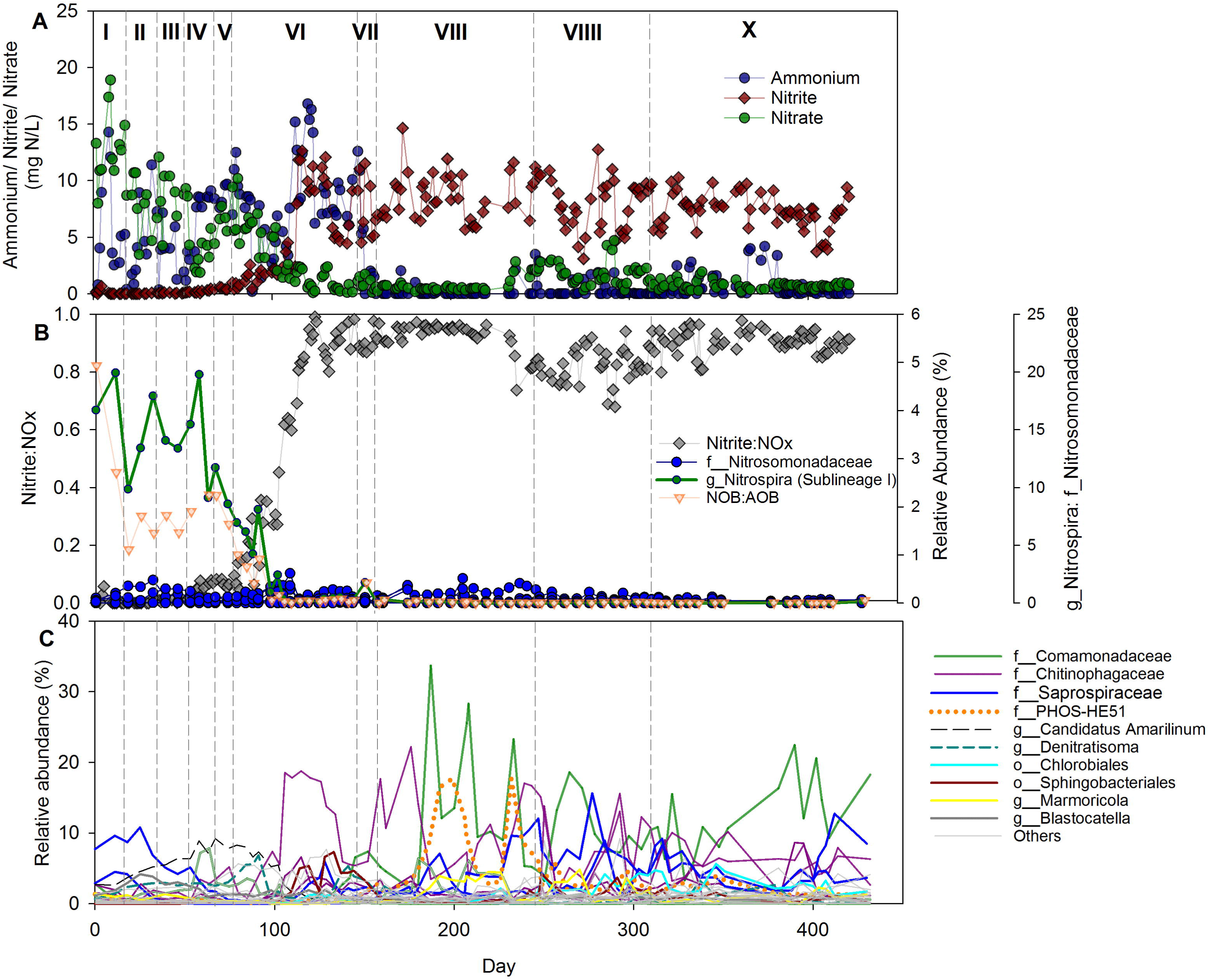
Effluent ammonium, nitrate and nitrite concentrations, (B) relative abundance of detectable ammonia oxidizing bacteria (f_*Nitrosomonadaceae*) and nitrite oxidizing bacteria (g_*Nitrospira*-sublineage I) and the corresponding nitrite:NOx ratio, and (C) relative abundance of all other detectable taxa in the sludge of the lab-scale partial nitritation(PN)/ nitritation(N)-denitritation(DN) sequencing batch reactor as a function of time. In (C) all detected taxa are displayed but only the top ten most abundant annotated taxa in all three samples are listed. Dashed lines indicate changes made to operational conditions summarized in Table 2.

In addition to *Nitrospira*, two OTUs annotated to the genera *Candidatus* Amarilinum and Blastocatella decreased in relative abundance with increasing DO concentration. The disappearance of *Nitrospira* also coincided with the proliferation of multiple heterotrophic taxa belonging to the families Chitinophagaceae, Saprospiraceae, Comamonadaceae and PHOS-HE51 (Figure 2C), which could be a consequence of increased availability of nitrite or organic carbon sources or both. These taxa displayed cyclical dynamics whereby the increase in relative abundance of one OTU was at the expense of another, indicative of competition for a limiting resource (Figure 2C). The changes in relative abundances were particularly apparent when the SRT was decreased to 8 days in experimental phase VIII, with a *Comamonadaceae* OTU increasing periodically in relative abundance up to approximately 34%. However, most other heterotrophic bacteria were present at relative abundances of <1% throughout all experimental phases.

In contrast to the effect on the NOB population in the activated sludge, we observed that ammonium oxidation was not compromised across a wide range of oxygen concentrations in the lab-scale reactor. While multiple AOB OTUs were detectable during the various experimental phases, different OTUs seemed to predominate at different DO set points, in line with the correlation between diversity and functional stability (Yachi and Loreau 1999). In addition, the AOB community appeared to be resilient with higher similarity between samples collected on day 56 (phase IV) and day 350 (phase X) than on day 212 (phase VIII) (Figure 1), indicating that the AOB community returned to a stage close to its original composition when the DO concentration was changed back to low concentrations after a long period of high oxygenation (Figures 1 and 2). Thus, increasing the oxygen set point in a NOB-rich sludge dominated by *Nitrospira* sublineage I resulted in inhibition of nitrite oxidation and eventual wash out of nitrite oxidizers from the system, consistent with the observation in the full-scale treatment plants that *Nitrospira* has a preference for lower oxygen concentrations compared to the coexisting AOB.

### Nitrifying activity in response to varying DO set points

To further understand the competitive dynamics between AOB and NOB in Plant 3 sludge for oxygen, the ammonia and nitrite oxidizing activities were characterized across a wide range of DO concentrations (Figure 3). Comparable nitrite oxidizing activities were observed in batch experiments in the presence of nitrite only (i.e. NOB activity test) and when both nitrite and ammonia were supplied (i.e. simultaneous AOB and NOB activity tests) at DO concentrations from 0.2 to 6.5 mg O_2_/L (Figure 4). In both NOB activity test and simultaneous AOB and NOB activity tests on Plant 3 sludge, the maximum specific nitrite oxidizing activity of approximately 6.0 mg N/h/g VSS was attained at a DO concentration of 1.5 mg O_2_/L. In contrast, a lower maximum specific ammonium oxidizing activity of 7.1 mg N/h/g VSS was observed in the simultaneous AOB and NOB batch activity tests compared to 9.3 mg N/h/g VSS in the AOB-only batch activity tests (i.e. ammonium only) (Figure 4). In the lab-scale reactor, following the wash out of the NOB, a much higher maximum ammonium oxidation rate of approximately 12.2 mg N/h/g VSS was achieved, suggesting that AOB activity in the sludge was significantly impacted when NOB activity was simultaneously occurring, whereas the NOB activity was not affected by concurrent oxygen consumption by AOB. The maximum AOB activities in all cases were higher than those of NOB and were attained at a DO concentration of 2.5 mg O_2_/L (Figure 4). However, in the simultaneous AOB and NOB batch activity tests at DO concentrations <1.5 mg O_2_/L, the nitrite oxidation rates were higher than the ammonia oxidation rates (Figure 4), further supporting that *Nitrospira* in Plant 3 sludge have lower relative *K*s for oxygen compared to *Nitrosomonas*-like AOB. The comparable nitrite oxidizing activity beyond DO concentration of 1.5 mg O_2_/L (up to 6.5 mg O_2_/L) also suggests that NOB are able to cope with the relatively high oxygen concentration and that the wash out of NOB observed in the long-term lab-scale study was from the imbalance in their ability to compete for oxygen with AOB when DO concentration was increased.

**Figure 3.**
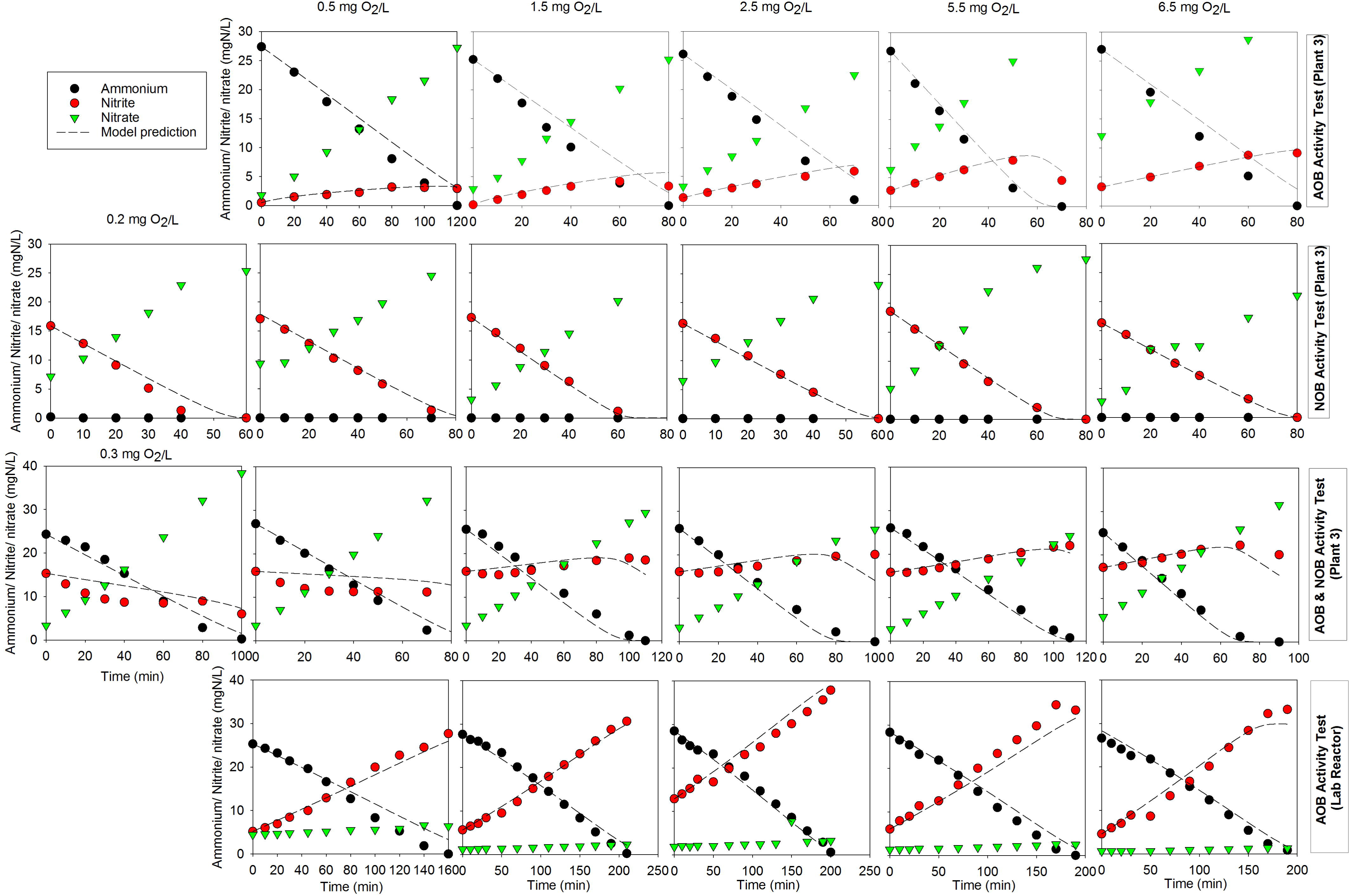
Experimentally observed and model-based ammonia oxidising and/or nitrite oxidising activity at varying dissolved oxygen concentrations in the ammonia oxidising bacteria (AOB) activity test (first row), nitrite oxidising bacteria (NOB) activity test (second row), and simultaneous AOB and NOB (third row) activity test with Plant 3 activated sludge and in the AOB activity test using sludge from the lab-scale reactor (fourth row). Each condition was tested in triplicate.

**Figure 4.**
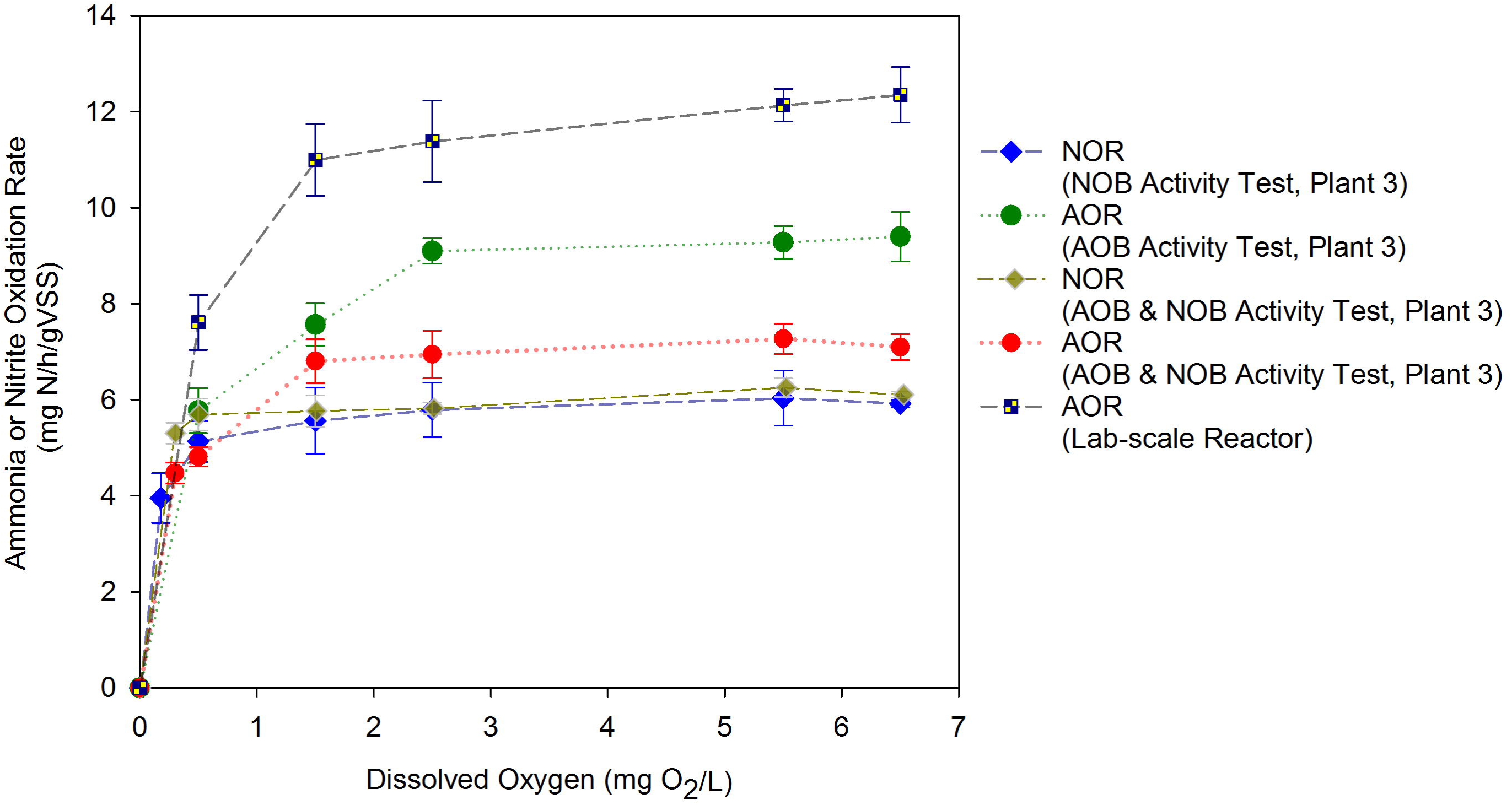
The effect of dissolved oxygen concentration on the average specific nitrite or ammonia oxidation rate (NOR or AOR) of activated sludge from Plant 3 in the nitrite oxidising bacteria (NOB) batch tests, ammonia oxidising bacteria (AOB) batch tests and the concurrent AOB and NOB batch tests and of sludge from lab-scale reactor in the AOB batch tests.

### Microbial kinetics of AOB and NOB

The various batch experiments were then modelled according to a Monod-based model for AOB and NOB (Wiesmann 1994) and the apparent *K*s_(app)_ and *µ*_max_ of the AOB and NOB were determined in batch tests (Figure 3). The good agreement between model predictions and measured results supported the validity of the estimated kinetic parameters (i.e., maximum growth rates and oxygen saturation constants of AOB and NOB) to describe the competitive dynamics between AOB and NOB at varying DO levels. The oxygen *Ks_(app)_* for AOB in Plant 3 sludge was estimated to be 0.30 ± 0.03 mg O_2_/L, higher than that of NOB with an estimated value of 0.09 ± 0.02 mg O_2_/L (Table S3, Supporting Information). In addition, the estimated *µ*_max_ of 0.126 ± 0.003 h^−1^ of AOB was almost ten times that of NOB, at 0.0128 ± 0.0003 h^−1^ (Table S3, Supporting Information). The estimated kinetic parameters suggest that *Nitrosomonas*-like AOB will outgrow *Nitrospira* in Plant 3 sludge when oxygen supply is high, whereas *Nitrospira* will predominate under oxygen limitation conditions; this could explain their high relative abundance in Plants 2 and 3. In addition, using the estimated kinetic parameters, the model was able to reproduce the batch experiment results with the lab-scale sludge, which further supports that the washout of *Nitrospira* was a result of operation at high DO.

### Physical properties of AOB and NOB microcolonies

To determine whether the lower *Ks_(app)_* of the NOB compared to AOB was an artifact of microcolony size, three dimensional FISH imaging was carried out on Plant 3 sludge to visualize the AOB and NOB colonies followed by image processing (Figure 5). Both AOB and NOB form cell clusters and reside in close proximity to each other. Image analysis revealed high variability in the volume and surface area of both AOB and NOB microcolonies (Figure S1, Supporting information). Cluster sizes are therefore not uniformly distributed in the sludge. However, while the total surface area of the AOB clusters were significantly larger compared to that of the NOB clusters (*p value = 0.0272*), the NOB clusters consistently occupied a higher volume than the AOB clusters (*p value = 0.0018*). Thus, despite inconsistent cluster sizes, the AOB microcolonies had significantly higher surface area to volume ratios than did NOB microcolonies (*p value < 0.0001*) and displayed far higher morphological variability. AOB mcrocolonies were less regular in appearance compared to those of NOB, which tended to be more spherical (Figures 5 and 6). The irregularity in shape resulted in conflicting maximum and minimum diameter of the microcolonies indicating that microcolony description based on size is not sufficient when describing substrate diffusion (Figure S2, Supporting Information).

**Figure 5.**
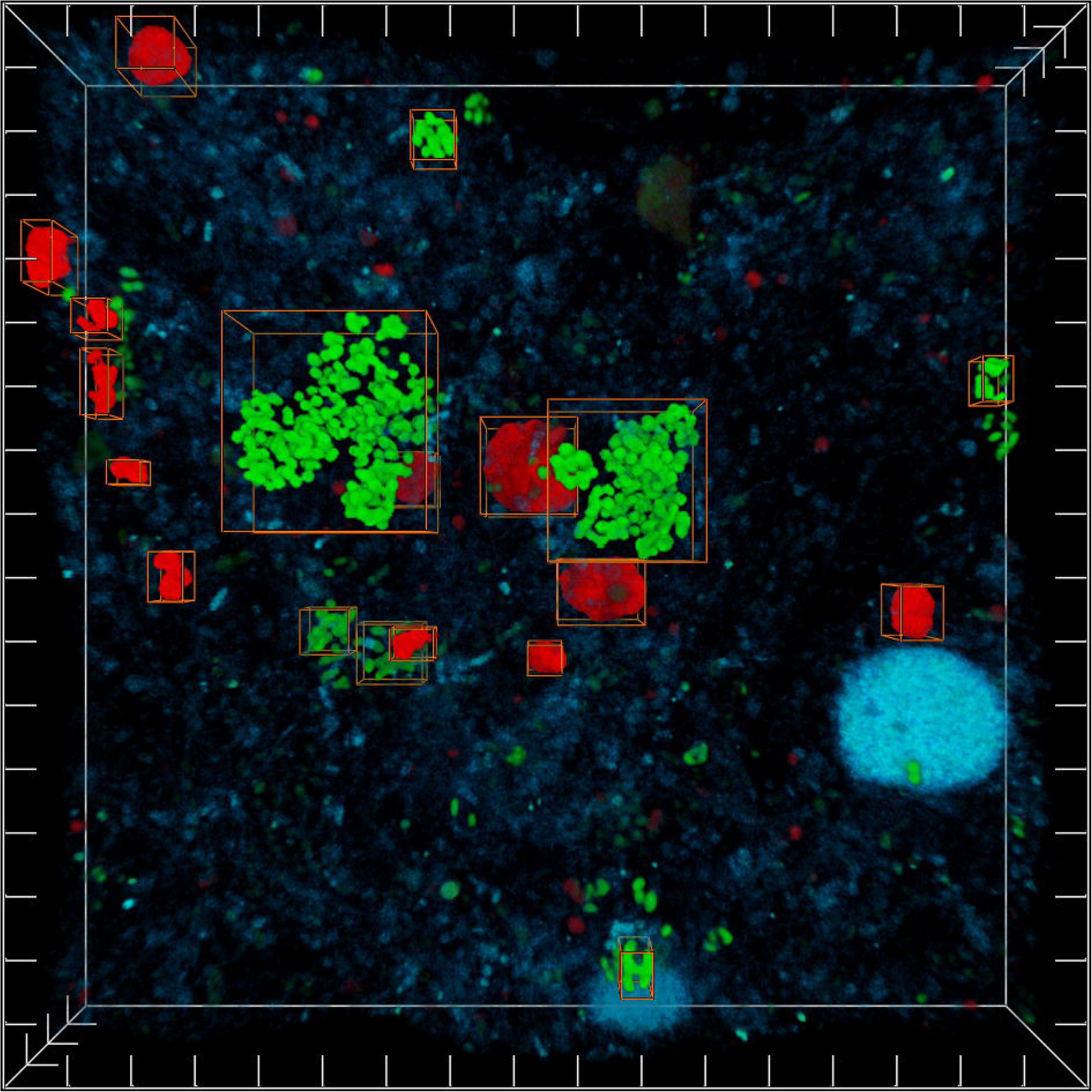
Ammonia oxidising bacteria (probes NSO1225 & NSO190 in green) and nitrite oxidising bacteria (probe Ntspa662 in red) colonies detected by fluorescence in situ hybridisation (FISH) in Plant 3 sludge. Boxes indicate microcolonies identified based on threshold described in materials and methods under further analyses. Ticks on all axes are uniformly distributed at 5 µm. All other detected bacteria are shown in blue (probes EUB338, EUB338-II and EUB338-III).

**Figure 6.**
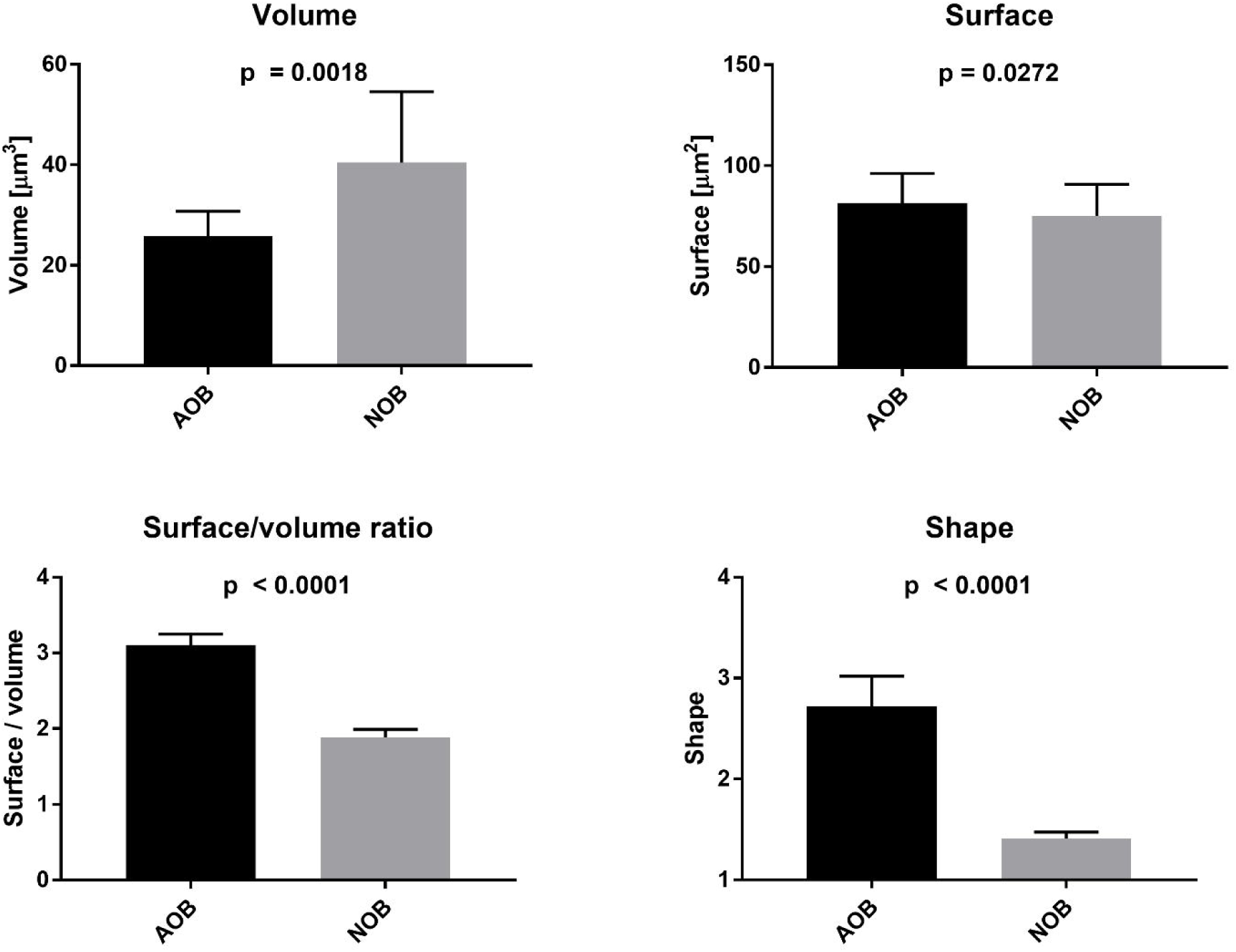
Physical properties of ammonia oxidising bacteria (AOB) and nitrite oxidising bacteria (NOB) clusters identified by fluorescence in situ hybridisation (FISH) with an example shown in Figure 4. Data represent the median with 95% confidence interval. P values are listed for unpaired nonparametric Mann-Whitney U test. The value for shape denotes the sphericity of the clusters with the value 1 being round.

## Discussion

Dissolved oxygen (DO) concentration is is a key factor regulating biogeochemical cycling in natural environments (Falkowski et al 2008) and also commonly used in wastewater treatment processes to shift microbial community dynamics towards operationally favored populations. In some instances this is obvious, such as when selecting for aerobes over anaerobes. In others, however, the application of DO as a tool to affect community composition and specific populations becomes more nuanced, such as selecting between two aerobes on the basis of different oxygen affinities. One example is selecting for AOB over NOB, which is attractive as a complement to Anammox in order to reduce operating costs. However, understanding the competition dynamics between AOB and NOB for oxygen is confounded by a combination of physical and biological factors acting at different scales, all the way from micro- to macro-scale (Arnaldos et al 2015, Picioreanu et al 2016). In this study, we resolved physicochemical and biological aspects of AOB/NOB interactions in full-scale activated sludge systems by integrating kinetic, microbial community and microcolony structure characterization along with field sampling and short term and long term lab-scale experimentation. Contrary to the paradigm that lower DO concentrations eliminate NOB in nitrogen rich side stream systems (Lackner et al 2014), we observed in this study into activated sludge systems receiving relatively low nitrogen loadings that the opposite was true.

Based on 16S rRNA gene amplicon sequencing, the NOB community composition was highly enriched compared to AOB within the same system, with a single *Nitrospira* sublineage I OTU dominating across different treatment plants in Singapore. When subjected to perturbations in oxygenation levels (Figure 2), *Nitrospira* numbers decreased in relative abundance with increasing DO concentration and were eventually washed out from the lab-scale system altogether (Figure 2). The nitrite oxidizing activity did not recover and sustained nitrite accumulation was observed even at low DO concentrations. The inability to reverse the nitrite oxidizing activity at lower DO concentration demonstrates that high DO operation may only be required for a short period of time to suppress NOB activity, after which oxygen levels can be adjusted depending on process requirement. This provides operational flexibility when combining partial nitritation with Anammox to achieve nitrogen removal, whereby a one-stage partial nitritation-Anammox system will require low oxygen levels to prevent inhibition of Anammox bacteria, whereas higher oxygen levels can be applied for a two-stage partial nitritation-Anammox system to maximize conversion rates. The prevalence of the same *Nitrospira* OTU across different treatment plants also suggests that under wastewater conditions in Singapore nitritation may be consistently achieved through high DO selection.

NOB are a highly diverse functional group that possesses fundamental ecophysiological diversity mainly stemming from differences in affinity to nitrite and energetic yield depending on the localization of the enzyme nitrite oxidoreductase (NXR) (Daims et al 2016). Environmental conditions such as DO concentration (Park and Noguera 2008), temperature (Alawi et al 2009, Daims et al 2001b, Siripong and Rittmann 2007) and pH (Hüpeden et al 2016) have also been shown to drive niche differentiation of NOB. The absence of OTUs from the other NOB genera in this study is not unexpected given the lack of temporal fluctuations in the plants and the year round stable wastewater temperature of 30 °C, which favours *Nitrospira* over *Nitrotoga* (Alawi et al 2009), and the relatively diluted ammonium content with a continuous flow configuration that limits the nitrite production rate by AOB. Such conditions favour the selection of *Nitrospira* taxa that possess a periplasmic NXR with a high nitrite affinity, allowing them to adapt to low nitrite concentrations (Lücker et al 2010, Schramm et al 1999) as opposed to *Nitrobacter* that have higher nitrite conversion rates but lower nitrite affinity (Kim and Kim 2006). On the contrary, excess ammonium availability will result in dynamic changes in ammonium profiles that potentially provide ecological niches for the co-occurrence of multiple AOB strains (Daims et al 2001b, Ke et al 2013). Mass balance analysis revealed a significantly higher proportion of NOB compared to AOB in Plants 2 and 3, suggesting that the growth of *Nitrospira* was not exclusively dependant on autotrophic nitrite oxidation. Indeed, mixotrophic growth has been shown for *Ca.* Nitrospira defluvii (Spieck et al 2006) and *Nitrospira marina* (Watson et al 1986). The *Ca.* N. defluvii genome also contains genes that encode pathways for the transport, oxidation, and assimilation of simple organic compounds (Lücker et al 2010). In the lab-scale reactor, the increase in a number of heterotrophic taxa after *Nitrospira* was washed out from the system suggests that they occupy the same ecological niche. The antagonistic relationships between OTUs annotated to Chitinophagaceae, Saprospiraceae and Comamonadaceae (Figure 2C) are more likely to be due to competition for organic carbon than nitrite given that nitrite was always in excess when *Nitrospira* had been inhibited. In the case of Plant 1 where an anaerobic zone is a plant design feature to maximise organic carbon uptake by heterotrophic bacteria, AOB were more abundant than NOB, further suggesting potential antagonistic interactions between *Nitrospira* and heterotrophs in activated sludge. However, the higher operational DO concentration and longer SRT may also have had an impact on the observed higher proportion of AOB to NOB. Nevertheless, nitrite did not accumulate in any of the investigated plants, indicating that it was consumed immediately either by NOB or by heterotrophic bacteria.

While 16S rRNA gene amplicon sequencing may not be able to provide subspecies level resolution for *Nitrospira* (Gruber-Dorninger et al 2015), the decline of the detected *Nitrospira* OTU in the long-term lab-scale experiment suggests that multiple species under sublineage I, if present, are not well adapted to high oxygen concentration. This was also reflected in the *Ks_(app)_* that was determined to be three-fold lower than that of AOB and also in the significantly lower estimated *µ*_max_, indicating that *Nitrospira* will thrive under low oxygen conditions. In addition, batch experiments showed that the AOB activity was significantly affected by the simultaneous oxygen consumption by NOB, in contrast to model predictions by Picioreanu *et al*. (2016). The *Ca.* N. defluvii genome shows the presence of the reductive tricarboxylic acid (rTCA) cycle, the anaerobic cobalamin biosynthesis pathway, and the lack of genes for protection against reactive oxygen species (ROS) normally present in most aerobic organisms, suggesting *Nitrospira* is of anaerobic or microaerophilic origin (Lücker et al 2010). In fact, comparative genomics revealed an evolutionary link between *Nitrospira* and anammox organisms (Lücker et al 2010). Both AOB and NOB are known to grow in microcolonies with near spherical shape (Daims et al 2001b, Mobarry et al 1996, Vejmelkova et al 2012, Wagner et al 1995). The growth of *Nitrospira* in aggregates has been proposed to potentially offer additional protection to cope with oxidative stress (Lücker et al 2010).

Our study shows that AOB and NOB exhibited microcolonies of distinct morphology when subjected to the same level of oxygenation in Plant 3 of 1.2 ± 0.2 mg O_2_/L (Figures 5 and 6). *Nitrospira* formed tightly packed and dense colonies with a lower surface area to volume ratio that limits mass transfer of oxygen in agreement with kinetic properties determined in this study and their preference for lower oxygen concentrations. In contrast, *Nitrosomonas*-like AOB formed open porous aggregates with a high surface area to volume ratio that maximises diffusional mass transfer. Porous colonies are expected to yield lower *Ks_(app)_* values (i.e., higher affinity), whereas dense colonies would lead to the reverse (Martins et al 2004). Yet, the estimated *Ks_(app)_* for NOB is still significantly lower than that of AOB, suggesting that the intrinsic oxygen affinity constant (i.e., not affected by diffusion) of NOB could be even lower than the estimated value in this study. While AOB are generally thought to form larger colonies than NOB based on the measurement of colony diameter (Coskuner et al 2005, Manser et al 2005, Picioreanu et al 2016), we show that colony characterization solely based on diameter may lead to skewed conclusions given the inconsistency between the maximum and minimum diameter of non-spherical colonies (Figure S2, Supporting Information). Therefore, our results are contrary to the contention of Picioreanu *et al*. (2016) that observation of a lower oxygen affinity for NOB could be the consequence of the difference in microcolony size.

Collectively, the findings suggest that nitrifiers may regulate microcolony structure formation depending on their intrinsic physiological and kinetic properties and environmental conditions. Such capability may assist them to survive under a broader range of environmental conditions and interact and coexist with partners that are adapted to distinct environments. For NOB, the lower surface area to volume ratio would impair oxygen capture, which in agreement with Lücker et al. (2010), could protect *Nitrospira* against oxidative stress allowing them to still perform nitrite oxidation at high oxygen concentrations up to 6.5 mg O_2_/L. In contrast, the porous microcolonies formed by the AOB in Plant 3 suggest that oxygen was limiting but such microcolony formation would allow maximum oxygen transfer to sustain ammonia oxidizing activity. Therefore, stable and active populations of *Nitrosomonas*–like *AOB* and *Nitrospira* can be maintained in domestic wastewater treatment plants between low and intermediate DO set points but the partnership can still be destabilised with a high operating DO as demonstrated in this study. Further investigation is required to understand whether microcolony structure formation is regulated by external stimuli or is a consequence of environmental selection for specific strains with a defined morphotype. The regulation of microcolony structure may also be a survival strategy for anaerobes to persist in suboxic to oxic environments.

## Acknowledgements

This research was supported by the Singapore National Research Foundation and Ministry of Education under the Research Centre of Excellence Programme, by a program grant from the National Research Foundation (NRF), project number 1301-IRIS-59, and the National Medical Research Council (NMRC/CBRG/0086/2015). We thank Mr. Larry Liew and staff from PUB, Singapore’s National Water Agency for performing weekly collection of primary effluent. Dr. Bing-Jie Ni acknowledges the support of Australian Research Council Future Fellowship FT160100195. We thank Dr. Kimberly Kline and Dr. Per Halkjær Nielsen for reviewing the manuscript.

