## Supplementary Materials for "Apparent oxygen half saturation constant for nitrifiers: genus specific, inherent physiological property, or artefact of colony morphology?"

(*Supplementary Information*)

Yingyu Law[1*], Artur Matysik[1], Xueming Chen[2], Sara Swa Thi[1], Thi Quynh Ngoc Nguyen[1], Guang Lei Qiu[1], Gayathri Natarajan[1], Rohan B.H. Williams[3], Bing-Jie Ni[4], Thomas William Seviour[1], Stefan Wuertz[5]*

[1] Singapore Centre for Environmental Life Sciences Engineering, Nanyang Technological University, Singapore 637551, Singapore

[2] Process and Systems Engineering Center (PROSYS), Department of Chemical and Biochemical Engineering, Technical University of Denmark, 2800 Kgs Lyngby, Denmark

[3] Singapore Centre for Environmental Life Sciences Engineering, National University of Singapore, Singapore 119077, Singapore

[4] Centre for Technology in Water and Wastewater, School of Civil and Environmental Engineering, University of Technology Sydney, Sydney, NSW 2007, Australia

[5] School of Civil and Environmental Engineering, Nanyang Technological University, Singapore.

Table S1. Composition of the primary effluent augmented with nitrite (values are average and standard deviation measured throughout the experiment).

| Parameter | Concentration |
| --- | --- |
| Total Kjeldahl nitrogen (mg N/L) | 39.4 ± 11.7 |
| Total Phosphorus (mg P/L) | 7.1 ± 1.1 |
| Ammonium (mg N/L) | 35.8 ± 4.4 |
| Nitrite (mg N/L) | 70.1 ± 10.3 |
| Phosphate (mg P/L) | 4.1 ± 1.7 |
| Total COD (mg/L) | 260 ± 40 |
| Total Alkalinity (mg/L $CaCO_3$) | 160 ± 25 |
| Acetate (mg/L) | 27.5 ± 13.1 |
| Propionate (mg/L) | 4.2 ± 2.5 |
| Butyrate (mg/L) | 0.3 ± 0.6 |

Table S2. Stoichiometric Matrix for the Model

| Process | $S_{NH4}$ N | $S_{NO2}$ N | $S_{NO3}$ N | $S_{O2}$ $O_2$ | $X_{AOB}$ COD | $X_{NOB}$ COD | $X_I$ COD | Kinetic rate expression |
|---|---|---|---|---|---|---|---|---|
| 1. Growth of AOB | $-i_{NBM}-\frac{1}{Y_{AOB}}$ | $\frac{1}{Y_{AOB}}$ | | $-\frac{3.43-Y_{AOB}}{Y_{AOB}}$ | 1 | | | $\mu_{AOB}\frac{S_{NH4}}{S_{NH4}+K_{NH4}^{AOB}}\frac{S_{O2}}{S_{O2}+K_{O2}^{AOB}}X_{AOB}$ |
| 2. Decay of AOB | $i_{NBM}-i_{NXI}*f_I$ | | | | $-1$ | | $f_I$ | $b_{AOB}X_{AOB}$ |
| 3. Growth of NOB | $-i_{NBM}$ | $-\frac{1}{Y_{NOB}}$ | $\frac{1}{Y_{NOB}}$ | $-\frac{1.14-Y_{NOB}}{Y_{NOB}}$ | | 1 | | $\mu_{NOB}\frac{S_{NO2}}{S_{NO2}+K_{NO2}^{NOB}}\frac{S_{O2}}{S_{O2}+K_{O2}^{NOB}}X_{NOB}$ |
| 4. Decay of NOB | $i_{NBM}-i_{NXI}*f_I$ | | | | | $-1$ | $f_I$ | $b_{NOB}X_{NOB}$ |

Table S3. Stoichiometric and Kinetic Parameters of the Model

| Parameter | Definition | Value | Unit | Source |
|---|---|---|---|---|
| *Stoichiometric parameters* | | | | |
| $Y_{AOB}$ | Yield coefficient for AOB | 0.150 | g COD $g^{-1}$ N | Wiesmann 1994 |
| $Y_{NOB}$ | Yield coefficient for NOB | 0.041 | g COD $g^{-1}$ N | Wiesmann 1994 |
| $i_{NBM}$ | Nitrogen content of biomass | 0.07 | g N $g^{-1}$ COD | Henze et al. 2000 |
| $i_{NXI}$ | Nitrogen content of $X_I$ | 0.02 | g N $g^{-1}$ COD | Henze et al. 2000 |
| $f_I$ | Fraction of $X_I$ in biomass decay | 0.10 | g COD $g^{-1}$ COD | Henze et al. 2000 |
| *Ammonium oxidizing bacteria (AOB)* | | | | |
| $\mu_{AOB}$ | Maximum growth rate of AOB | 0.126±0.003 | $h^{-1}$ | This work |
| $b_{AOB}$ | Decay rate of AOB | 0.0054 | $h^{-1}$ | Wiesmann 1994 |
| $K_{NH4}^{AOB}$ | $S_{NH4}$ half saturation constant for AOB | 1.1 | g N $m^{-3}$ | Wiesmann 1994 |
| $K_{O2}^{AOB}$ | $S_{O2}$ half saturation constant for AOB | 0.30±0.03 | g $O_2$ $m^{-3}$ | This work |
| *Nitrite oxidizing bacteria (NOB)* | | | | |
| $\mu_{NOB}$ | Maximum growth rate of NOB | 0.0128±0.0003 | $h^{-1}$ | This work |
| $b_{NOB}$ | Decay rate of NOB | 0.0025 | $h^{-1}$ | Wiesmann 1994 |
| $K_{NO2}^{NOB}$ | $S_{NO2}$ affinity constant for NOB | 0.5 | g N $m^{-3}$ | Wiesmann 1994 |
| $K_{O2}^{NOB}$ | $S_{O2}$ affinity constant for NOB | 0.09±0.02 | g $O_2$ $m^{-3}$ | This work |

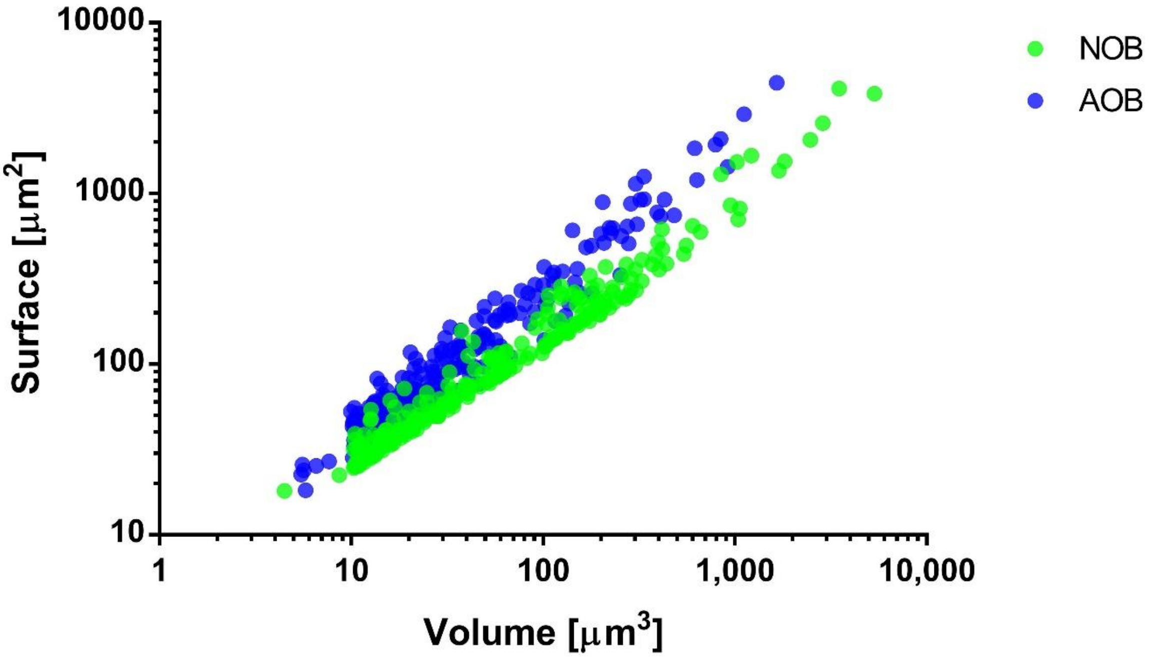


Figure S1 The distribution of the surface area and volume of ammonia oxidizing bacteria (AOB) and nitrite oxidizing bacteria (NOB) colonies in Plant 3 sludge identified by fluorescence in situ hybridisation (FISH).

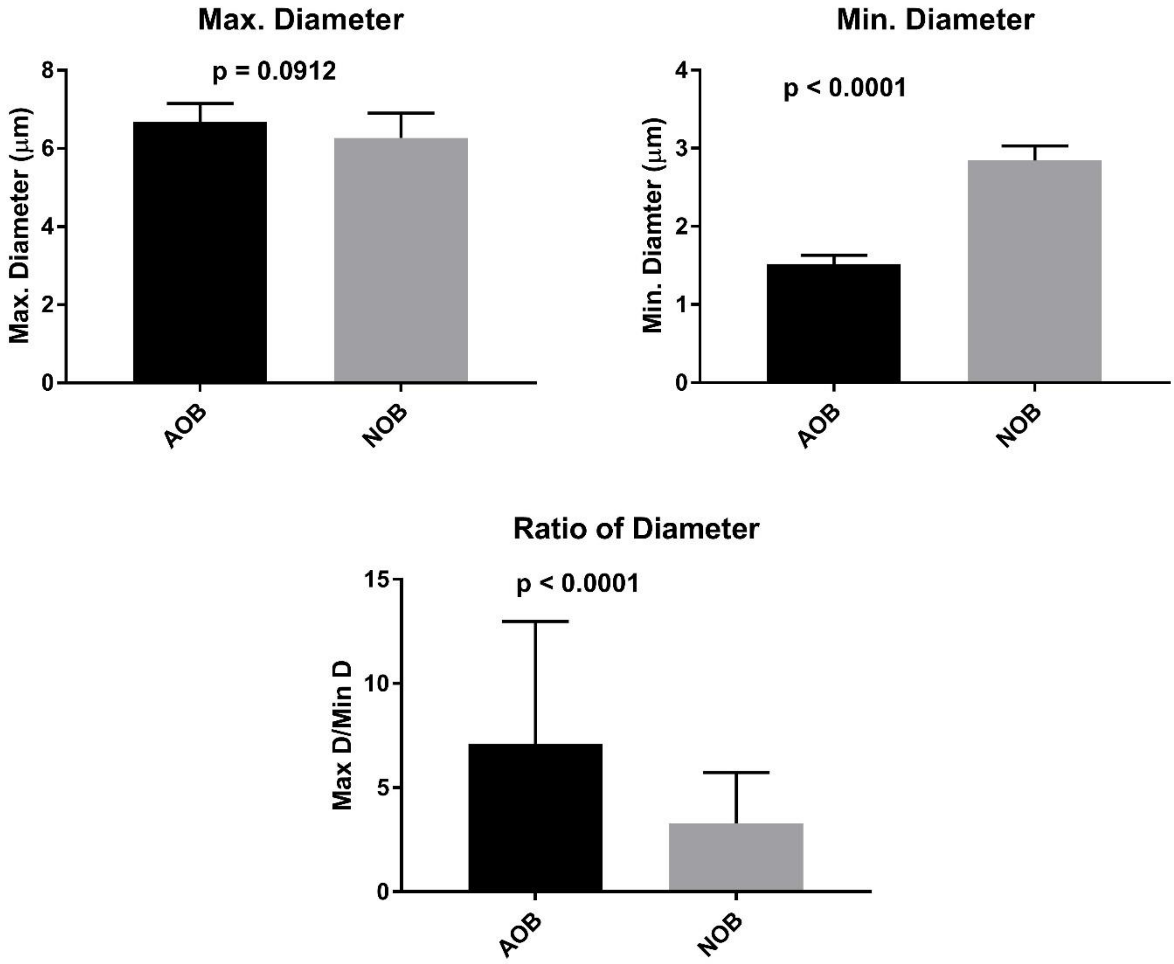


Figure S2 The maximum and minimum diameter of the ammonia oxidizing bacteria (AOB) and nitrite oxidizing bacteria (NOB) colonies in Plant 3 sludge identified by fluorescence in situ hybridisation (FISH). Data represents median with 95% confidence interval.
